# Envelope-Limited Chromatin Sheets (ELCS) Formation in The Nuclear Envelope of HL-60/S4 Cells

**DOI:** 10.64898/2026.02.23.707298

**Authors:** Ada L. Olins, Igor Prudovsky, Donald E. Olins

## Abstract

Envelope-Limited Chromatin Sheets (ELCS) can be induced in human promyelocytic HL-60/S4 cells by treatment with retinoic acid (RA). After 4 days, the differentiated granulocytes exhibit multilobed nuclei with outgrowths of the nuclear envelope (NE) and associated heterochromatin extending into the surrounding cytoplasm (ELCS). These fascinating structures reveal a periodic meshwork of 30 nm chromatin fibers, when viewed by Cryo-electron microscopy. Genetic and biochemical evidence indicates that RA increases the synthesis of Lamin B Receptor (LBR), which is a key enzyme for Cholesterol biosynthesis and is an essential bridge between the NE and peripheral heterochromatin. This article is in part a review of our microscopic data on the structure of ELCS, and in part a description of related transcription changes that result in the formation of ELCS. In addition, this article contains a structural and biochemical comparison of RA-induced granulocytes with phorbol ester (TPA) induced HL-60/S4 macrophages, which lack nuclear lobulation, do not form ELCS, and exhibit a reduction in LBR and Cholesterol biosynthesis. From our perspective, ELCS can be viewed as “fabric” outgrowths of the nuclear envelope, frequently connecting nuclear lobes and capable of sustaining the twisting and squeezing distortions imposed upon nuclear shape, as the granulocytes traverse narrow tissue channels.

## Introduction

Envelope-Limited Chromatin Sheets (ELCS) are an expansion of the interphase nuclear envelope, where the inner nuclear membrane (INM) and its affiliated layer of heterochromatin fibers are folded into a “sandwich” consisting of two apposed INM and their affiliated heterochromatin, exhibiting a symmetric arrangement. ELCS have been observed in numerous plants, animals and protozoa (Olins and Olins, 2009). H.G. Davies and co-workers studied ELCS early (Davies, 1968; Davies and Haynes, 1975; Davies et al., 1974; Davies and Small, 1968). They employed conventional “thin-section transmission electron microscopy (TEM)” (i.e., aldehyde fixation, dehydration, plastic embedding and sectioning, followed by Uranyl and Lead staining), yielding high quality electron micrographs. Davies and co-workers referred to the chromatin fibers of ELCS as “Superunit Threads” (Davies et al., 1974; Davies and Small, 1968). Molecular models of Superunit Threads remained ill-defined until after the discovery of the nucleosome, see review (Olins and Olins, 2003). In 1998, we observed that *in vitro* differentiation of a human myeloid cell line (HL-60/S4) with retinoic acid (RA) produced granulocytes with extensive ELCS, employing TEM examination (Olins et al., 1998). Our measurements of the distance between the apposed INMs agreed with those of Davies et al; i.e., ∼30 nm thickness (Figure 1). In subsequent collaborative studies (Eltsov et al., 2014; Xu et al., 2021) employing cryo-electron microscopy (Cryo-EM) to prevent sample shrinkage of HL-60/S4 granulocyte cells, it became clear that the distance between the two apposed INMs is ∼60 nm (Figure 2). Furthermore, Cryo-EM demonstrated that the thickness of each (hydrated) heterochromatin sheet is ∼30 nm; yielding two sheets of parallel 30 nm chromatin fibers forming a “criss-cross” pattern adjacent to the apposed INMs (Eltsov et al., 2014; Xu et al., 2021). The generally accepted chromatin fiber model became a 30 nm diameter fiber with a helical arrangement of nucleosomes. The present article attempts to understand the biochemical expansion of the interphase nuclear envelope and associated 30 nm heterochromatin fibers into the remarkably ordered ELCS formation.

**Figure 1.**
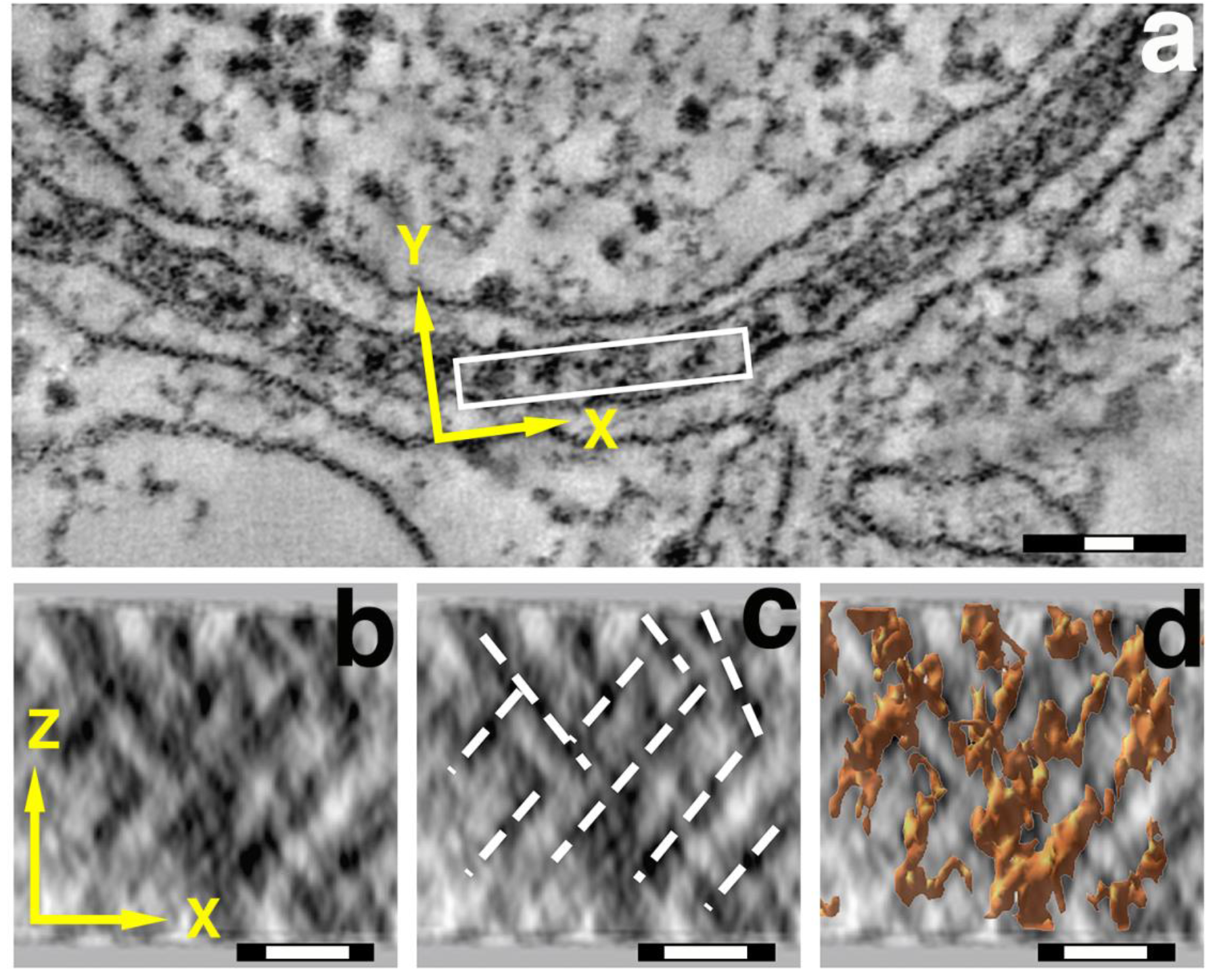
TEM images of ELCS. Top panel: **a**) end-views down the Z-axis of chromatin “Superunit Threads” between apposed inner nuclear membranes. Magnification bars: black 100 nm, white 30 nm. Bottom panels: **b, c, d**) Y-axis projections of the same ELCS region, illustrating the chromatin criss-cross pattern (**b**), the fiber pathways (**c**), and superimposed surface-rendering of “Superunit Threads” (**d**). Magnification bars: black 50 nm, white 30 nm. Images previously published (Eltsov et al., 2014).

**Figure 2.**
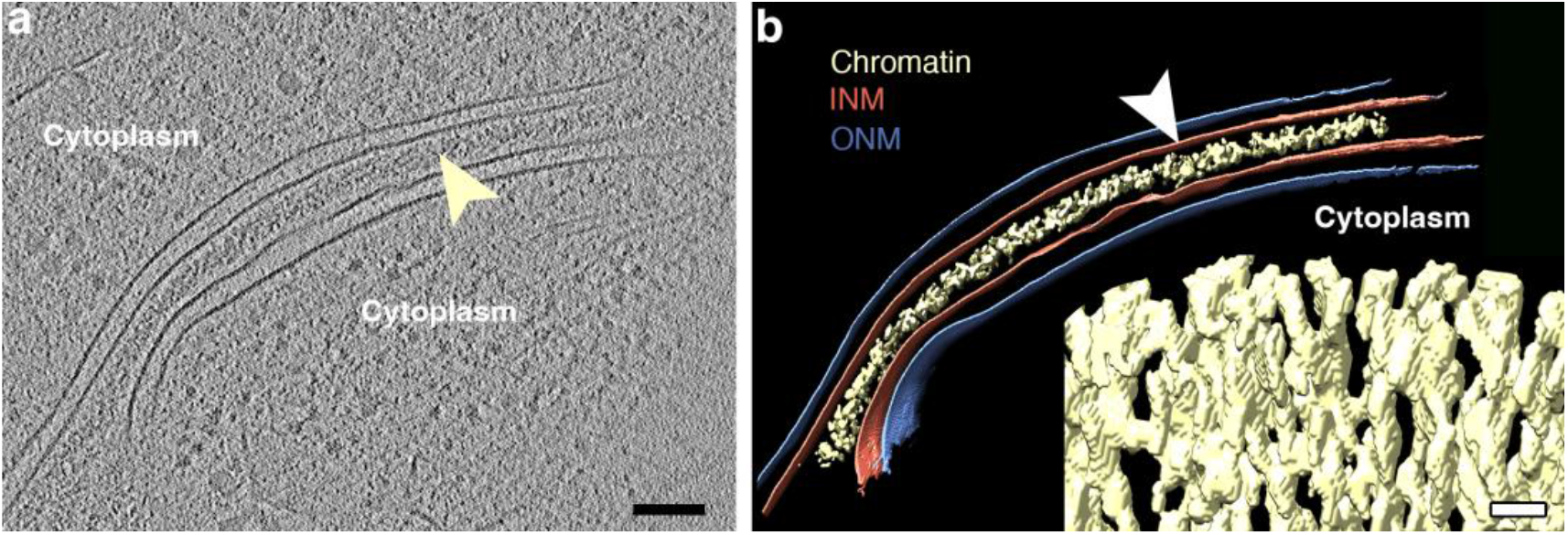
Cryo-EM image (**a**) and surface rendering (**b**) of ELCS substructure. In the left panel, the yellow arrowhead points to the chromatin affiliated with the apposed INMs. In the right panel, the white arrowhead points to the surface-rendered INM. The lower right corner insert of this panel displays a surface-rendering of the ELCS criss-cross chromatin pattern. The black magnification bar equals 100 nm and applies to panels **a & b.** The white magnification bar in the insert equals 20 nm. These images have been previously published (Xu et al., 2021).

## Results

### Electron microscopy of Nuclei in Several Different Cell States

Most of our nuclear structural studies have employed the derived HL-60/S4 cell line (Leung et al., 1992), which require less time to undergo cell differentiation than the parent HL-60 cell line. Many of our differentiated comparisons have been with RA-treated S4 cells (granulocytes) compared to untreated (0) S4 cells (Controls). Some comparisons have used phorbol ester (TPA) to differentiate S4 cells into macrophages. HL-60/S4 granulocytes grow in suspension; HL-60/S4 macrophages adhere to coverslip or glass slides (Mark Welch et al., 2017). Figure 3 presents TEM comparison images for 0, RA and TPA interphase nuclei.

**Figure 3.**
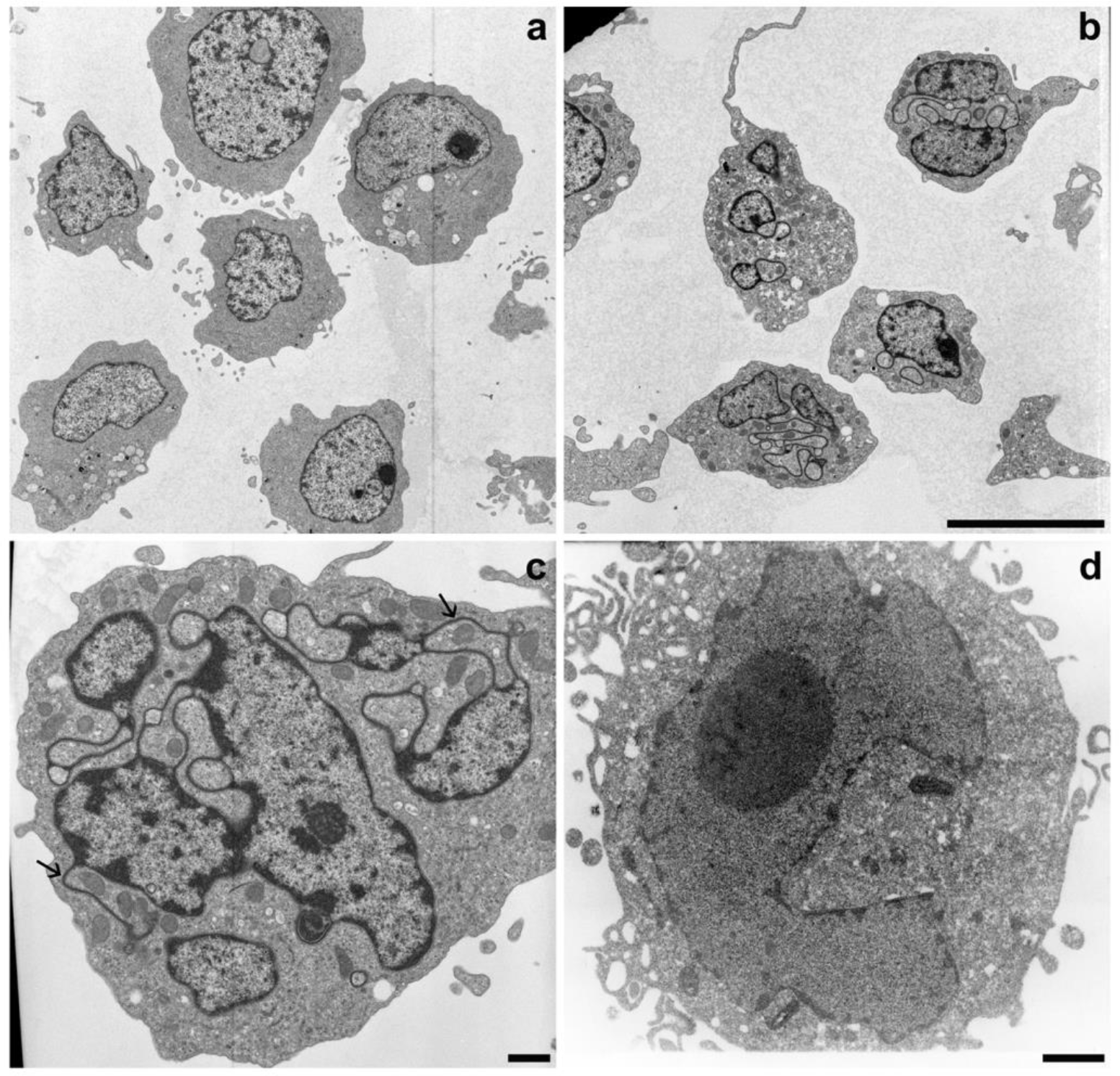
TEM images of 0, RA and TPA treated HL-60/S4 cells. Panel **a**: Undifferentiated S4 cells. Note the absence of nuclear lobulation and ELCS. Panel **b**: RA treated S4 cells. Note the presence of multilobed nuclei, frequently associated with ELCS. The magnification bar for both panels (**a** and **b**) equals 10 µm. Panel **c**: Higher magnification image of a RA treated S4 cell. Note the nuclear multilobulation with associated ELCS, many with microlobes. The two arrows point to apparent “nuclear pockets” with mitochondria trapped inside (Ghadially et al., 1985). Nuclear pockets are regions of cytoplasm surrounded by ELCS and/or lobules. Panel **d**: TPA treated S4 cell. Note the absence of nuclear lobulation and ELCS. The magnification bars for both panels **c** and **d** equal 1 µm.

### Composition of ELCS

Because it is not yet possible to biochemically purify ELCS fractions from contiguous nuclear lobes, it has been necessary to infer composition using cytochemical and immunochemical microscopy. Figure 4 displays TEM sections from two different RA treated S4 cells stained specifically for DNA, employing the Osmium Ammine-B (OA-B) method (Derenzini et al., 2014). OA-B stained ELCS can be clearly observed connected to adjacent stained nuclear lobes, which display thicker densely stained regions.

**Figure 4.**
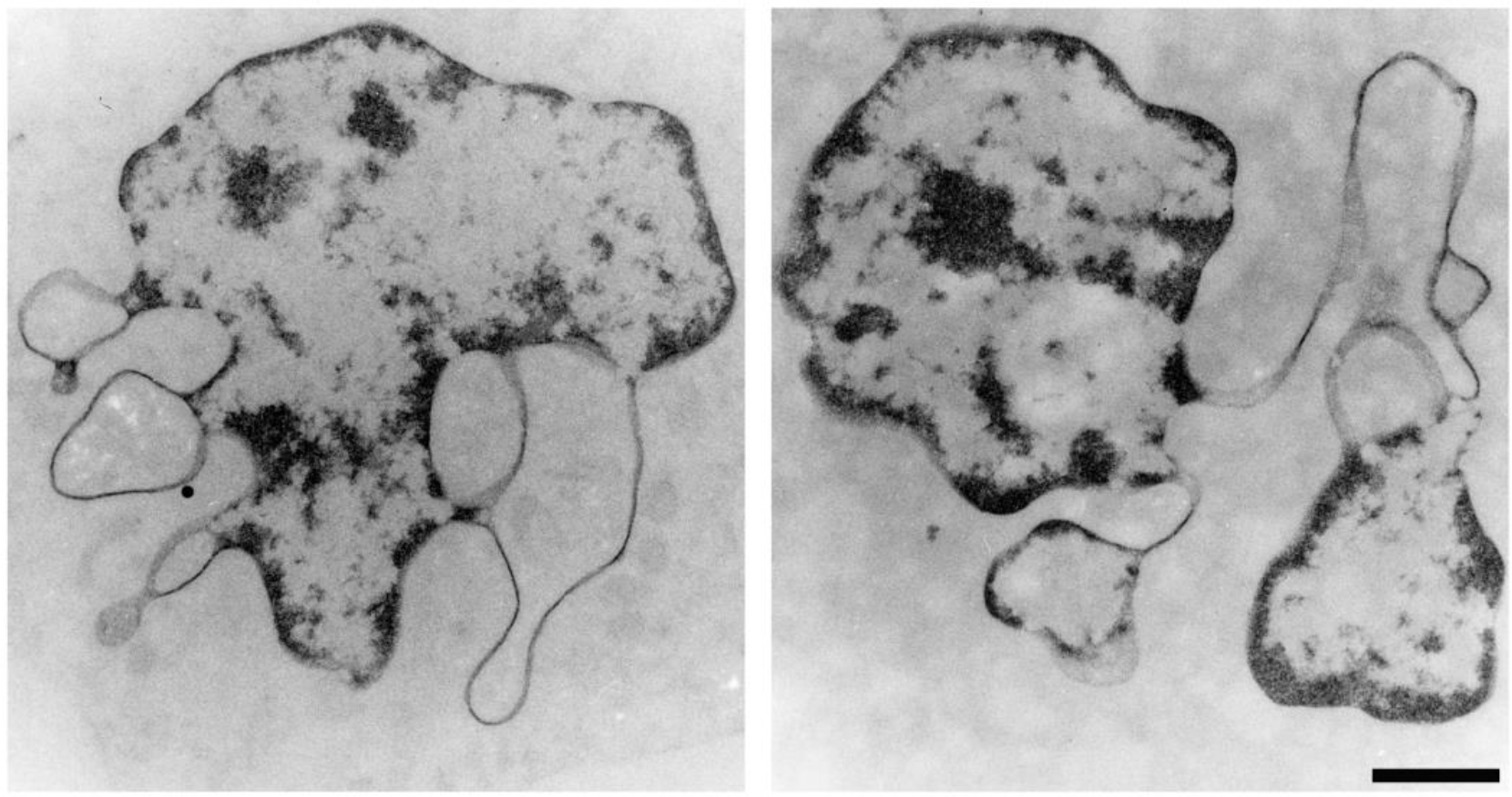
TEM images of RA differentiated HL-60/S4 granulocytes stained specifically for DNA, employing osmium-ammine B (Derenzini et al., 2014). The magnification bar for both panels equals 1 µm. This figure was initially published in (Olins et al., 1998).

Immuno-electron microscopy was employed to obtain evidence that a nucleosome epitope (“acidic patch”) is present within ELCS and adjacent chromatin regions of RA differentiated HL-60/S4 granulocytes (Figure 5). A mouse monoclonal antibody (PL2-6) has been demonstrated to bind at a junction of nucleosome histones (H2A and H2B). The bivalent and papain-derived monovalent versions of PL2-6 clearly react with the exposed nucleosome acidic patch “Epichromatin” epitope (Gould et al., 2017; Olins et al., 2011; Zhou et al., 2019).

**Figure 5.**
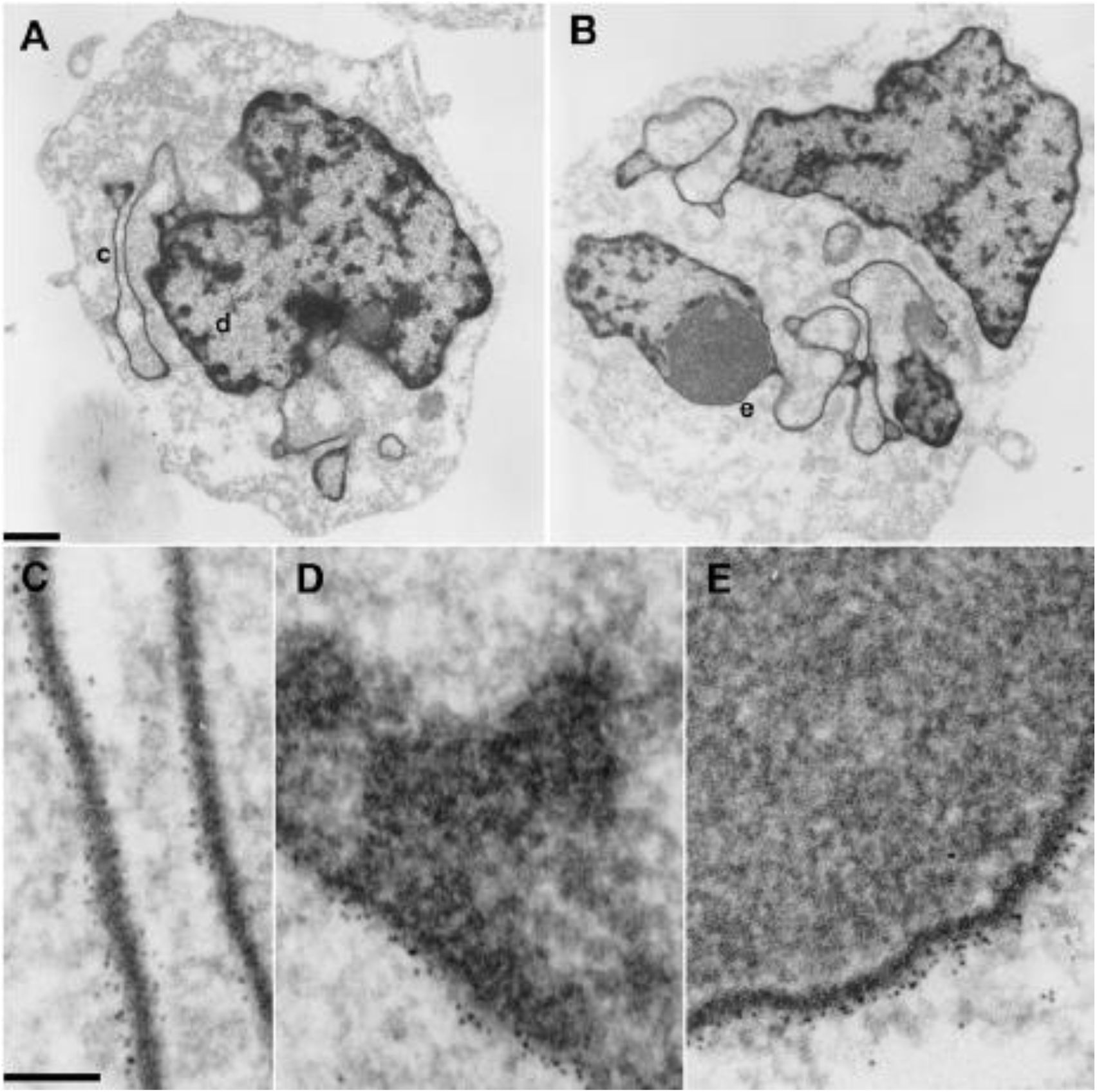
Immuno-TEM localization of the acidic patch epichromatin epitope within the nuclei of RA-treated S4 granulocytes. Panels A and B: Two different granulocytes with three substructures highlighted (**c**, ELCS; **d**, nuclear periphery; **e**, nuclear envelope adjacent to a nucleolus). The bottom three panels C, D and E display at higher magnification the immuno-gold particles bound to these three highlighted substructures. The top magnification bar is 1 µm; the bottom bar indicates 0.1 µm. Immuno-TEM localization of lamin B within ELCS has also been previously published (Olins et al., 1998).

Immunofluorescent confocal imaging studies (Figures 6 and 7) demonstrate that Lamin B Receptor (LBR) and heterochromatin protein 1 (CBX5) both stain nuclear envelope regions of HL-60/S4 granulocytes that are enriched with ELCS.

**Figure 6.**
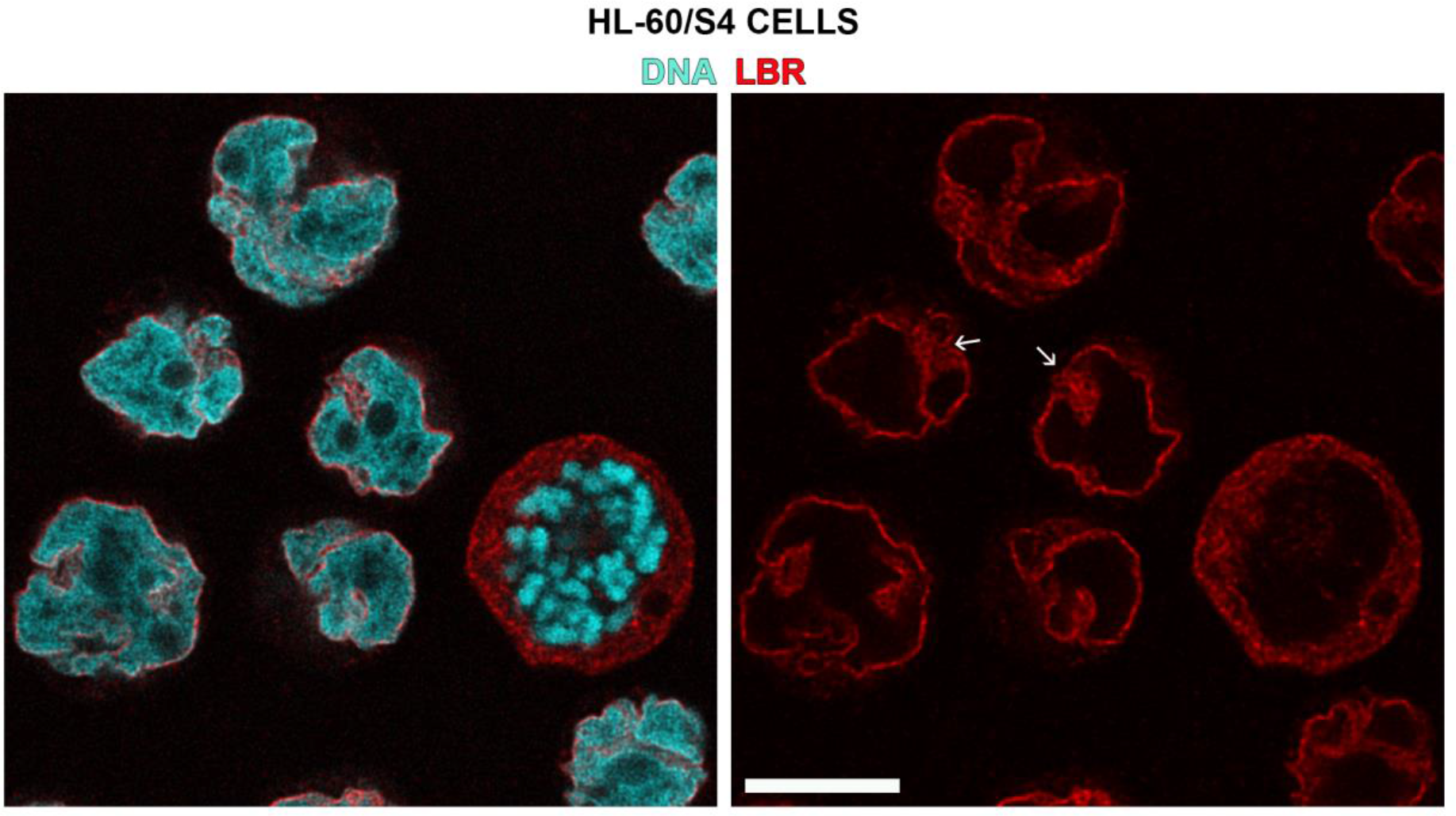
Immunofluorescent staining of RA-treated HL-60/S4 granulocytes (LBR, red; DNA, cyan). The small white arrows point to clusters of ELCS (between nuclear lobes) stained with Rabbit anti-LBR (Abcam ab32535, 1:500 dilution). Note that anti-LBR stains the nuclear envelope regions outside of ELCS. Note also, the accumulation of LBR (red) within the cytoplasm of a mitotic cell. The magnification bar for both panels equals 10 µm.

**Figure 7.**
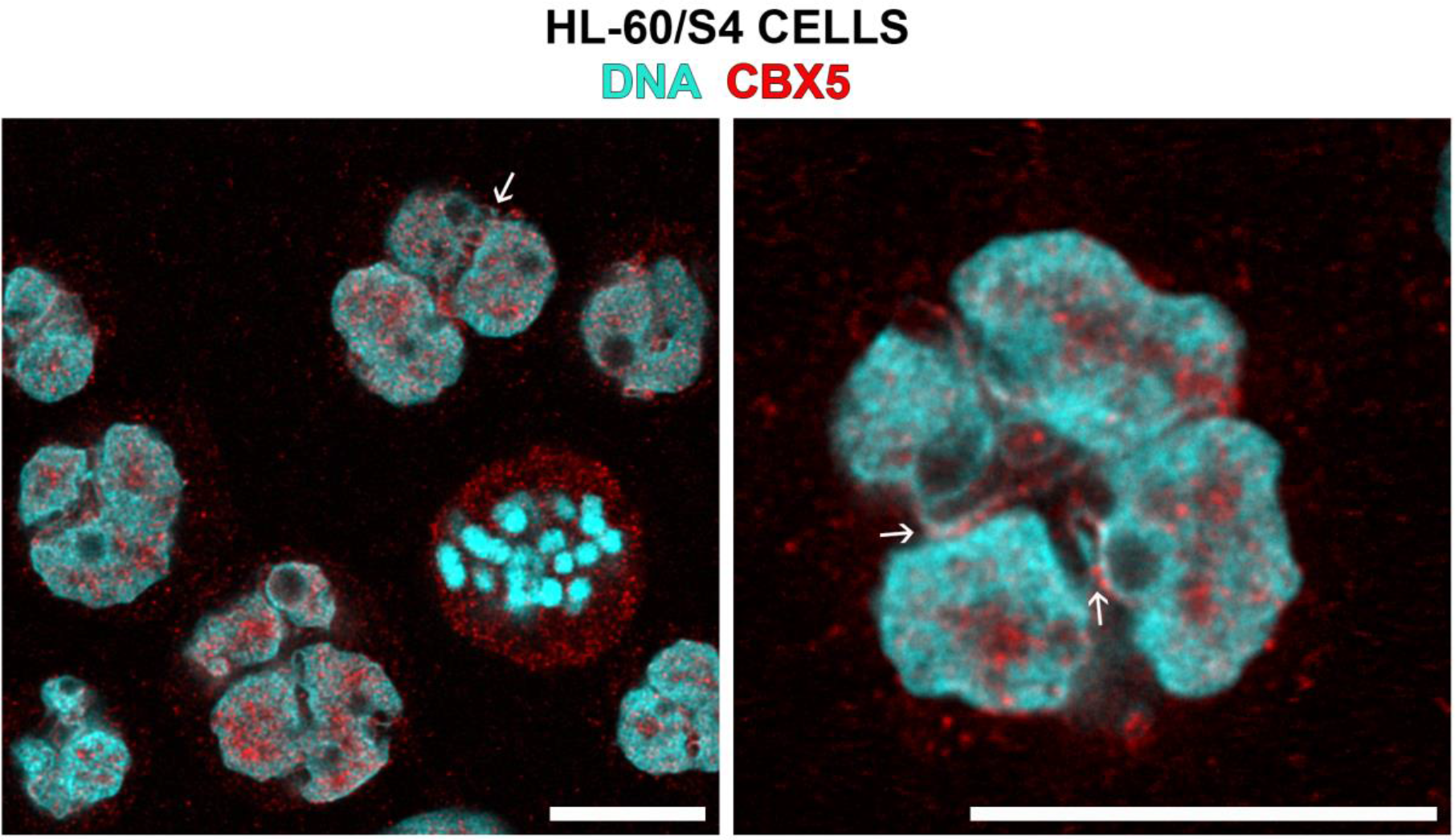
Immunofluorescent staining of RA-treated HL-60/S4 granulocytes (CBX5, red; DNA, cyan). The small white arrows point to ELCS (between nuclear lobes) stained with Goat anti-CBX5 (Abcam ab77256, 1:50 dilution). Note also, the accumulation of CBX5 (red) within the cytoplasm of a mitotic cell. The magnification bar for both panels equals 10 µm.

### Transcriptome Analyses: Searching for how HL-60/S4 cells can develop lobulation and ELCS after RA differentiation, but not after TPA differentiation

The goal of this portion of the Results is to speculate on what gene transcripts might be responsible for ELCS formation and growth during induced granulocyte differentiation. The multiplicity of ELCS following RA treatment requires significant increases in the biosynthesis of nuclear envelope membrane components. Employing morphometric analyses of nuclear envelope membrane areas in thin-sections of RA versus 0 cells indicated that NE area more than doubled (Olins et al., 1998).

Expansion of the NE membrane into lobes and ELCS clearly requires increased biosynthesis of lipids. Cholesterol is one of the most important lipids, even at its low concentration in the nuclear membrane and the endoplasmic reticulum (∼3-6% of total lipids) compared to the cell (plasma) membrane (∼30-50% of total lipids) (Niu and Balla, 2025; Ridsdale et al., 2006).

Cholesterol functions to stiffen nuclear membrane structure including formation of Lipid Rafts, while maintaining proper membrane fluidity (Nikolakaki et al., 2017; Shahoei and Nelson, 2019; Yang et al., 2016). Transcriptome analysis (Figure 8) presents a comparison of differential gene expression (DGE) for Cholesterol biosynthesis enzymes, comparing differentiated versus control cells (i.e., RA/0 and TPA/0). The two major Cholesterol biosynthesis pathways (Block & Kandutsch-Russell) both appear to operate in the HL-60/S4 cells. However, it is likely that the Kandutsch-Russell pathway predominates in the RA cells, but not the TPA cells. Note that the log_2_FC change of the DHCR7 differential gene expression is 1.05 for the RA/0 and −0.01 for the TPA/0.

**Figure 8.**
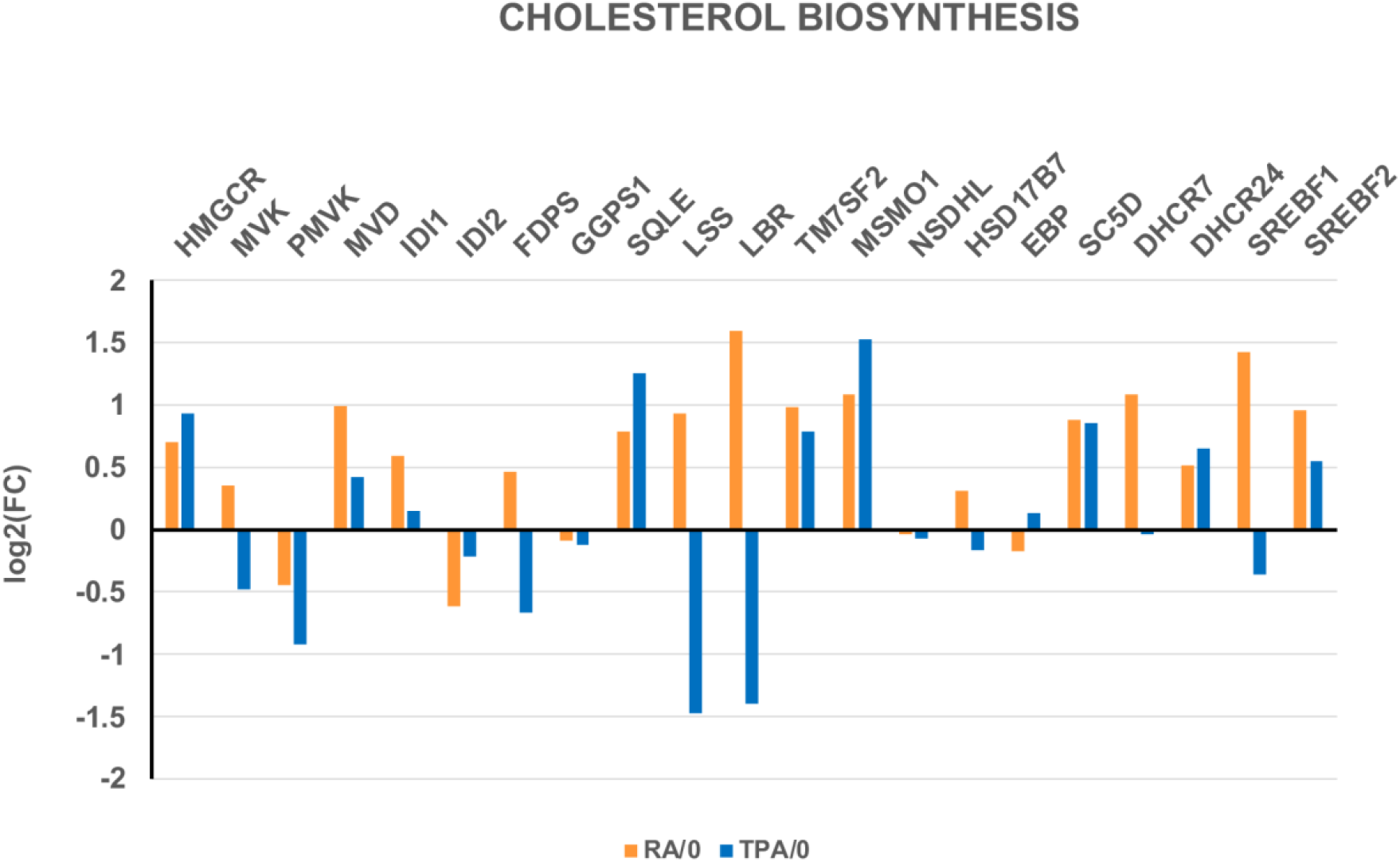
Comparison of differential gene expression (DGE) levels for both RA- and TPA-treated HL-60/S4 cells. Note that LBR is greatly increased by RA, and greatly reduced by TPA. In addition, note that DHCR7 (the final enzyme of the Kandutsch-Russell pathway) exhibits a significant increase in RA/0, but only a negligible negative change in TPA cells (Figure 8 and 9).

**Figure 9.**
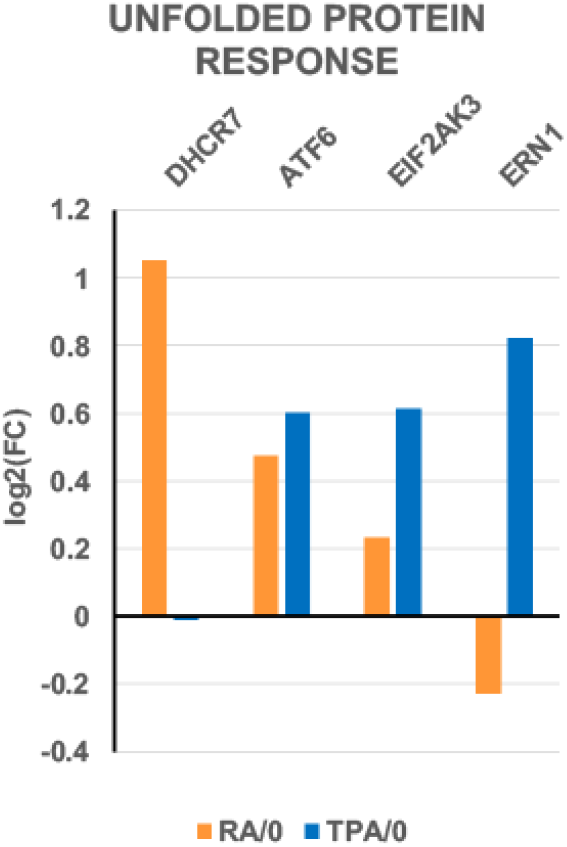
Unfolded Protein Response Normally, DHCR7 act as a protective enzyme by removing the toxic, proapoptotic 7-dehydrocholesterol (7-DHC). The miniscule change in expression of DHCR7 in TPA-treated HL-60/S4 cells, compared to the significant increase in RA-treated cells creates an important effect upon TPA cells: The “Unfolded Protein Response” is activated to battle “ER Stress”, by halting translation and by degrading misfolded nascent proteins. Figure 9 displays the significant increases in ATF6, EIF2AK3 and ERN1 transcription. These three primary transmembrane “sensor” proteins mediate the Unfolded Protein Response. It is interesting that “The Unfolded Protein Response” may be related to our prior observation that TPA-treated HL-60/S4 cells (but not RA-treated cells) are differentiated into Senescent Macrophages (Olins et al., 2024).

The DGE levels of nuclear envelope proteins (Figure 10) convey information about the structure and interactions within the interphase nuclei following treatment with RA and TPA. Figure 10 presents Log_2_FC plots of identical NE genes after RA and TPA differentiation. Perhaps the most interesting result in RA cells is the increased DGE of LBR, compared to its decrease in TPA cells. LBR performs multiple critical roles: including, sterol biosynthesis and acting as a nuclear structural protein (Olins et al., 2010b). Also striking in Figure 10 are the increased DGE for TPA-treated HL-60/S4 cells of LMNA, PLEC, VIM, SUN1, SYNE1 and SYNE2, compared to decreases or lesser increases in RA-induced granulocytes. Many of these DGE changes were noted in our earlier publication “The LINC-less Granulocyte Nucleus” (Olins et al., 2009). In that article, the authors hypothesized that the decrease of the LINC complex in RA induced granulocytes with lobulated nuclei serves to facilitate migration through tight tissue spaces. In other words, the granulocyte is a very malleable cell.

**Figure 10.**
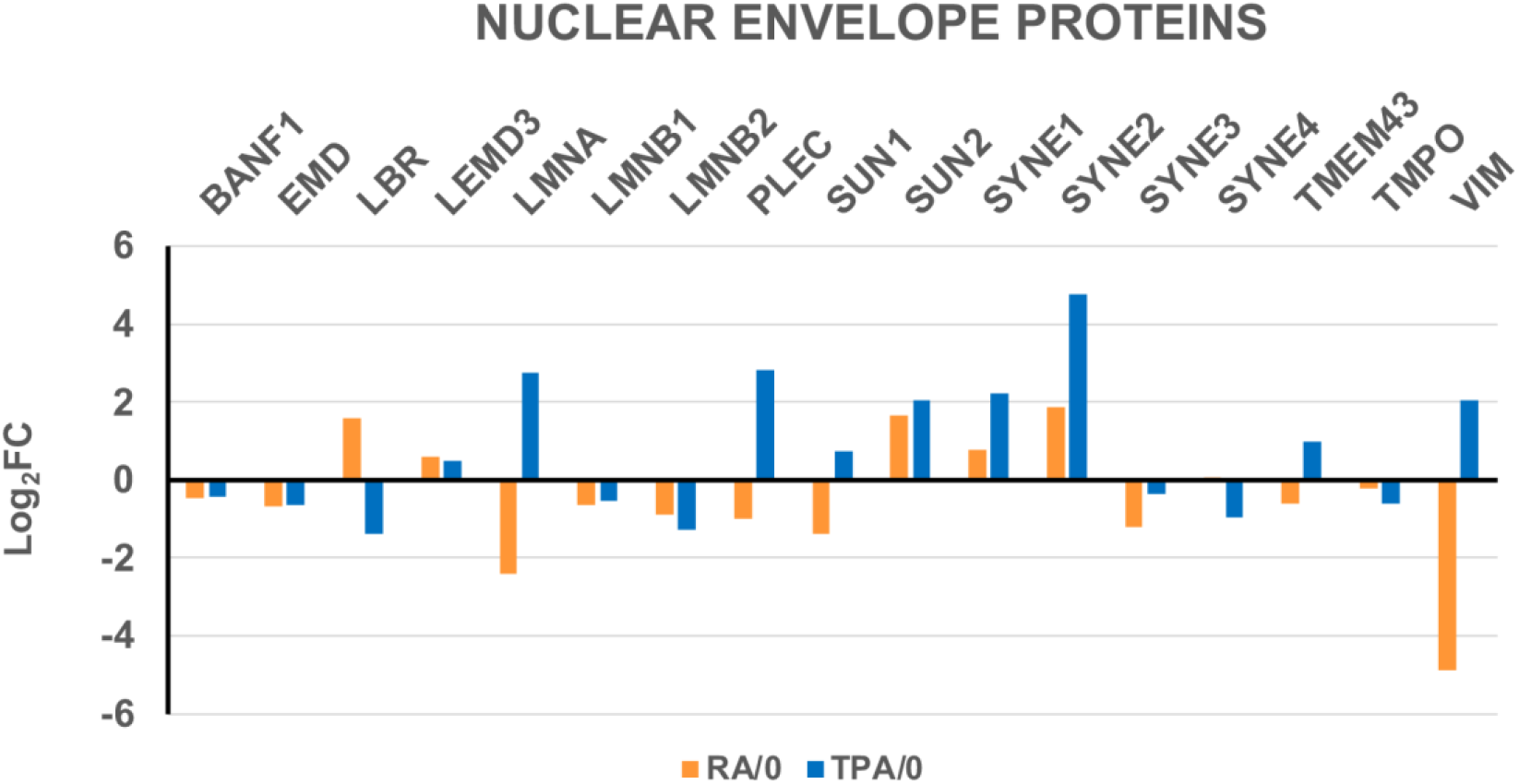
Comparison of the differential gene expression (DGE) levels for Nuclear Envelope Proteins in both RA- and TPA-treated HL-60/S4 cells.

### Functional Analyses

An alternative method to compare cholesterol biosynthesis after RA and TPA treatment is to apply Gene Set Enrichment Analysis (GSEA) to the transcription data. This method compares a “gene set” characteristic for a particular cellular phenotype to the experimental transcription data (in this case, RA/0 and TPA/0). A more complete description of this method can be found in an earlier publication from our laboratory (Olins et al., 2024). Figure 11 and Table 1 employ four different gene sets related to cholesterol synthesis, comparing the RA or TPA phenotypes to the undifferentiated cell state (0). For example, if most of a particular gene set is “bunched up” at the RA phenotype end of a plot, that particular GSEA function is enriched (NES>0) with transcriptionally-active genes. Low “Nominal p” values (i.e., close to 0.0) signify that the NES values are statistically significant; high values (i.e., close to 1.0) signify that the NES values are statistically not significant.

**Table 1.** GSEA plot parameters from Figure 11. Normalized Enrichment Score (NES) permits comparisons between different GSEA analyses, by accounting for variations in different gene set sizes. The results of these analyses clearly indicate that for all tested Cholesterol synthesis gene sets, the RA/0 NES values were greater than the TPA/0 NES values, and the Nominal p values were more significant for RA/0, compared to TPA/0.

| Gene Set | Phenotypes | NES | Nominal p | Genes |
| --- | --- | --- | --- | --- |
| REACTOME_CHOLESTEROL_BIOSYNTHESIS | RA/0 | 1.61 | 0.01 | 27 |
|  | TPA/0 | 0.61 | 0.98 |  |
| WP_CHOLESTEROL_METABOLISM | RA/0 | 1.81 | 0.0 | 72 |
|  | TPA/0 | 1.31 | 0.05 |  |
| REACTOME_CHOLESTEROL_BIOSYNTHESIS_VIA_DESMOSTEROL_BLOCH_PATHWAY | RA/0 | 1.43 | 0.06 | 10 |
|  | TPA/0 | 0.98 | 0.53 |  |
| GOBP_CHOLESTEROL_BIOSYNTHETIC_PROCESS_VIA_DESMOSTEROL | RA/0 | 1.53 | 0.02 | 19 |
|  | TPA/0 | 0.64 | 0.96 |  |

**Figure 11.**
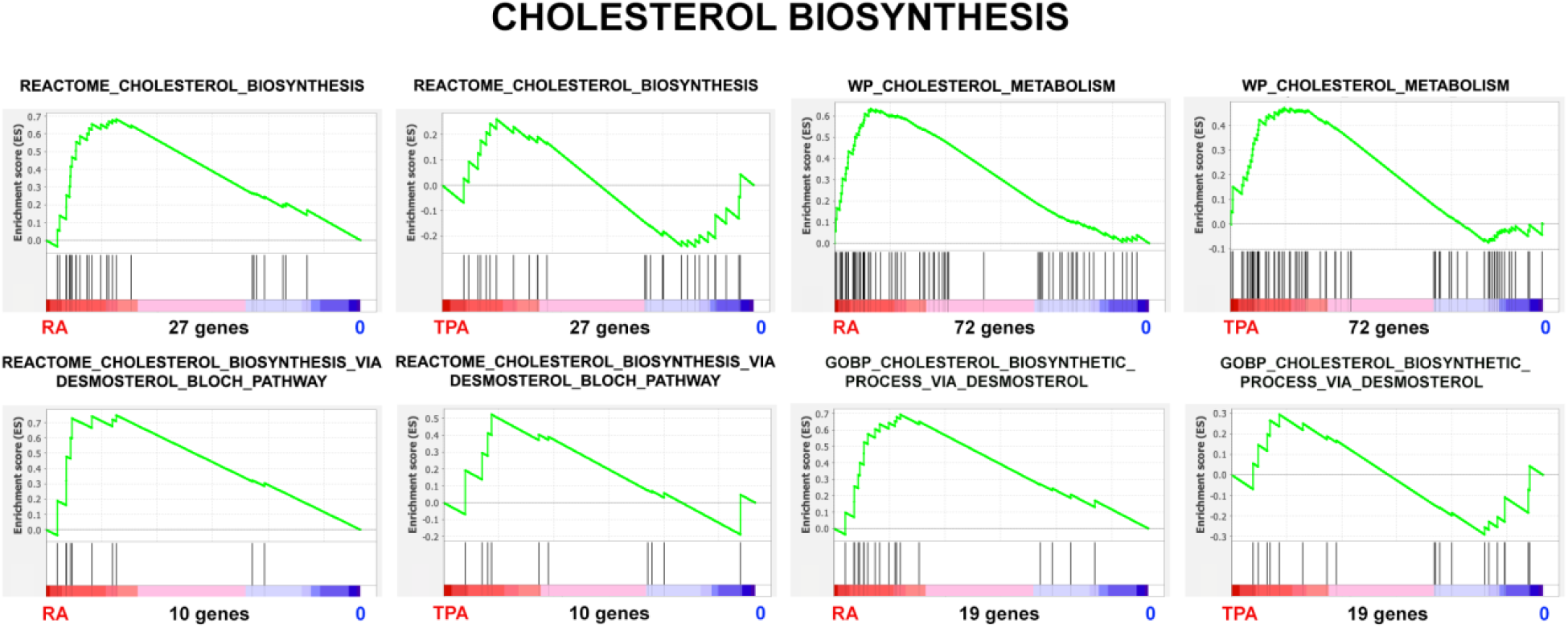
Cholesterol Biosynthesis. Gene Set Enrichment Plots of the phenotypes RA/0 and TPA/0. The magnitude of the Y-axes varies from plot-to-plot, reflecting the specific enrichment score (ES).

Each of these four GSEA Cholesterol gene sets has a “Leading Edge”; i.e., a subset of genes (“core group”) within a gene set that contributes most to the enrichment score. We compared these four Leading Edges (see Supplementary Materials Table S1) using Venn Analysis to determine what genes were common to all four leading edges. We determined that four genes (i.e., SC5D, MSMO1, LBR, DHCR7) were common to the four RA/0 Cholesterol leading edges. These four proteins (enzymes) were encountered earlier in Figure 8, illustrating their importance to Cholesterol biosynthesis. By contrast, analysis of the four TPA/0 Cholesterol leading edges included only three genes (i.e., SC5D, MSMO1, DHCR24). LBR transcription is down in TPA cells and therefore does not contribute to the leading edge. A different sterol double bond reductase, DHCR24 is in the leading edge for TPA/0 cells. It is interesting to note that DHCR24 is a part of the Bloch pathway for Cholesterol biosynthesis, while DHCR7 is a part of the of the Kandutsch-Russell pathway.

Glycerophospholipids constitute ∼65% of total nuclear lipids, compared to cholesterol ∼3-6 % of total nuclear lipids (Umair et al., 2025). Glycerolipids include glycerophospholipids and monoacyl, diacyl and triacylglycerol. All of these lipid categories appear to exhibit significantly increased NES, with TPA/0 being slighter greater than RA/0 (see Figure 12 and Table 2).

**Table 2.**
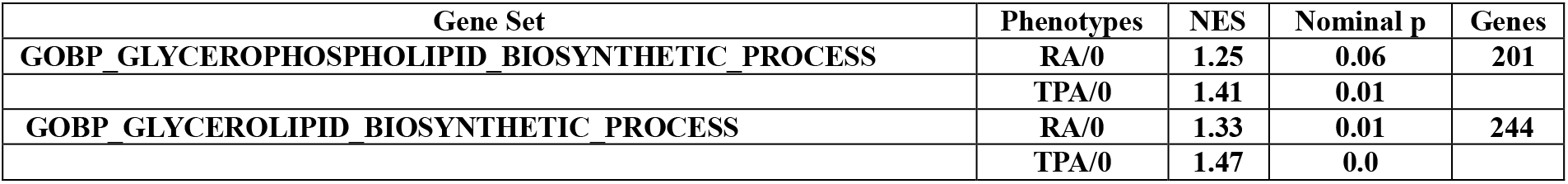
GSEA plot parameters from Figure 12. Normalized Enrichment Score (NES) permits comparisons between different GSEA analyses, by accounting for variations in different gene set sizes.

| Gene Set | Phenotypes | NES | Nominal p | Genes |
| --- | --- | --- | --- | --- |
| GOBP_GLYCEROPHOSPHOLIPID_BIOSYNTHETIC_PROCESS | RA/0 | 1.25 | 0.06 | 201 |
|  | TPA/0 | 1.41 | 0.01 |  |
| GOBP_GLYCEROLIPID_BIOSYNTHETIC_PROCESS | RA/0 | 1.33 | 0.01 | 244 |
|  | TPA/0 | 1.47 | 0.0 |  |

**Figure 12.**
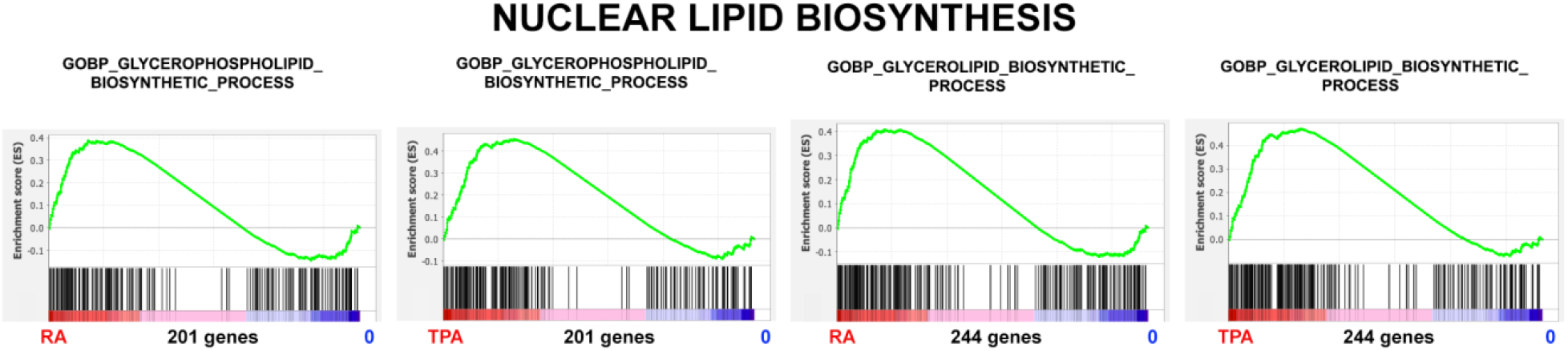
Glycerophospholipid and Glycerolipid Biosynthesis. Gene Set Enrichment Plots of the phenotypes RA versus 0, and TPA versus 0. The Y-axes varies from plot-to-plot, reflecting the specific enrichment score (ES).

**Figure 13.**
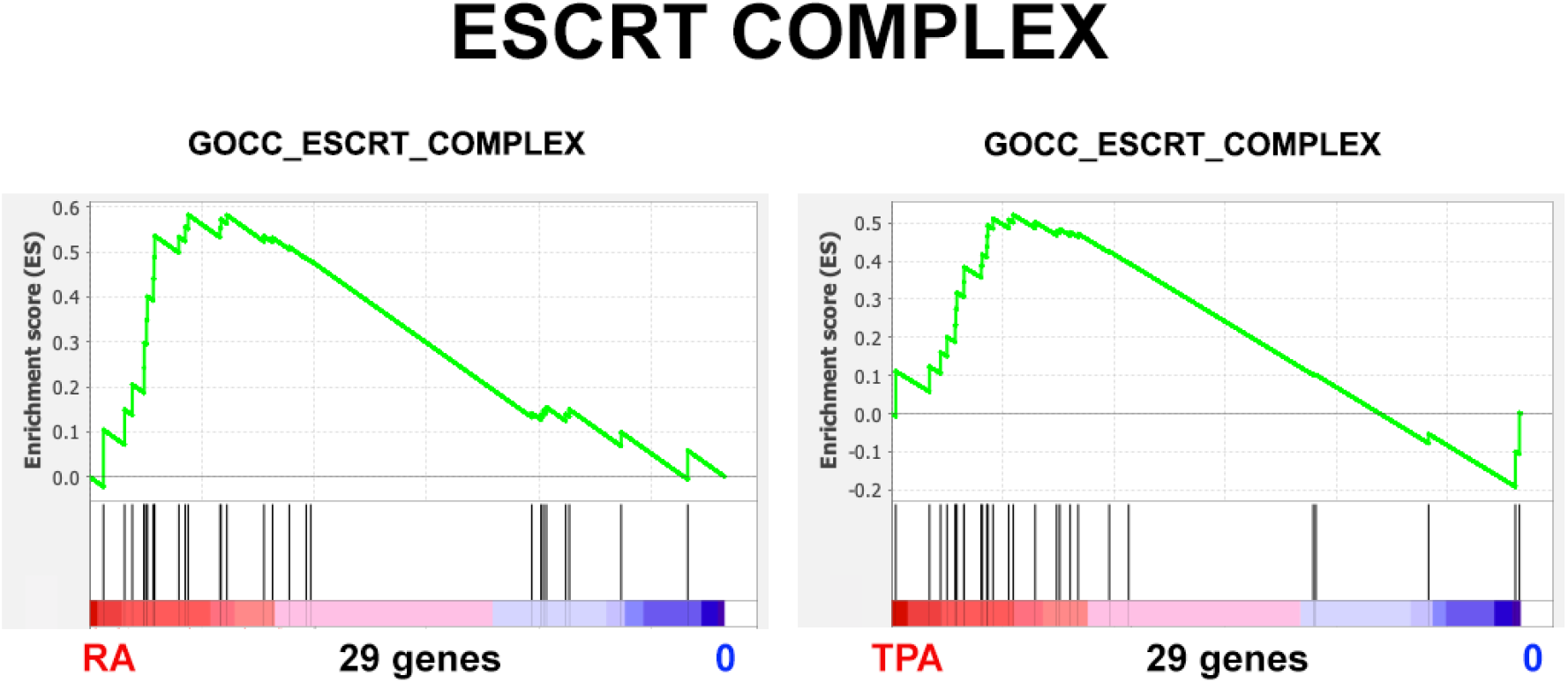
ESCRT Complex. Gene Set Enrichment Plots of the phenotypes RA/0, and TPA/0. The Y-axes vary from plot-to-plot, reflecting the specific enrichment score (ES).

We propose that the combination of Cholesterol and Glycerolipid biosynthesis favors the expansion and growth of the nuclear envelope in ELCS. The elevation of LBR in RA-differentiated cells promotes the formation of Lipid Rafts, due to the increase in localized Cholesterol biosynthesis. By contrast, the reduction of LBR in TPA-differentiated cells would disfavor the formation of Lipid Rafts (see Figures 8 and 10). LBR is “anchored” in the INM (Cheng et al., 2022) with heterogeneous mobility within the INM (Giannios et al., 2017). Lipid Rafts cluster LBR molecules which, in turn, tether heterochromatin to the INM of ELCS via LBR and CBX5 (Nikolakaki et al., 2017; Olins and Olins, 2009).

The ESCRT Complex can deliver endosomal vesicles containing lipids and proteins to the nuclear envelope (Hurley, 2015; Shankar et al., 2022; Wang et al., 2023). This machinery can repair or remodel the INM. Relative to the present study, the question is whether the ESCRT Complex can remodel the HL-60/S4 granulocyte and macrophage nuclear envelopes equally well. Examination of the plot parameters (Table 3) indicates that the RA-treated cells show a greater NES and a more significant Nominal p value than the TPA-treated cells. We suggest that the “regularity” of ELCS structure may make ESCRT remodeling an easier task.

**Table 3.** GSEA plot parameters from Figure 13. The higher NES values and lower Nominal p values of RA/0, compared to TPA/0, suggests that the ESCRT Complex may assist ELCS formation and expansion.

| Gene Set | Phenotypes | NES | Nominal p | Genes |
| --- | --- | --- | --- | --- |
| GOCC_ESCRT_COMPLEX | RA/0 | 1.42 | 0.04 | 29 |
|  | TPA/0 | 1.23 | 0.17 |  |

## Discussion

The major conclusion of this study is: The presence of adequately increased amounts of LBR per cell (Log_2_FC RA/0 equal to 1.56) is crucial for the formation and growth of Envelope-Limited Chromatin Sheets (ELCS) in retinoic acid (RA) treated granulocyte HL-60/S4 cells. The decrease of LBR (Log_2_FC TPA/0 equal to −1.37) in phorbol ester (TPA) treated macrophage HL-60/S4 cells probably insures that ELCS and nuclear lobes are not formed. We suggest that the amount of LBR per cell is the principal factor behind these nuclear shape changes. LBR has, at least, two functions: 1) a major enzyme for Cholesterol biosynthesis; 2) a structural protein, connecting heterochromatin and lamins to the Nuclear Envelope (Nikolakaki et al., 2017; Olins et al., 2010b).

How do ELCS form and grow? We suggest that higher-order chromatin fibers (i.e., “30 nm fibers”) affiliated with the INM (see Figures 1 and 2) continue to increase in amount following the RA-induced cessation of cell division. Histone H1-10 (alias, H1FX) continues to be synthesized, since it is a Replication-Independent H1 variant gene. We expect it to stabilize the higher-order chromatin structure. The Log_2_FC of H1-10 for RA/0 (0.76) and TPA/0 (−0.51) argues that 30 nm chromatin fibers can continue to form after RA differentiation, but do not increase after TPA treatment. Increased LBR, situated within the INM via its C-terminal transmembrane helices, binds to heterochromatic 30 nm chromatin fibers via its N-terminal Tudor Domain and its CBX5 binding site (Olins et al., 2010b). We further suggest that newly synthesized lipids (e.g., Glycerolipids) join the INM from the endoplasmic reticulum and the ESCRT Complex, establishing additional Lipid Rafts with additional Cholesterol, which would then bind nascent LBR, continuing the expansion of the Nuclear Envelope area and the enrichment of ELCS. Presumably, TPA induced macrophages do not exhibit nuclear lobulation and ELCS because of the deficiency of LBR, histone H1-10 and Lipid Rafts.

The centrality of LBR to nuclear lobulation and the formation of ELCS in HL-60/S4 granulocytes is further substantiated by additional observations/publications: 1) knockdown of LBR in HL-60/sh1 cells eliminates lobulation and ELCS formation following RA treatment (Mark Welch et al., 2024; Olins et al., 2010a; Olins et al., 2025); 2) human individuals with an increased number of LBR genes in their genomes exhibit hyperlobulation of their neutrophils (Gravemann et al., 2010).

Regulation of LBR mRNA biosynthesis following treatment of cells with RA is well understood. All-trans retinoic acid binds to a nuclear receptor complex (RAR+RXR), which subsequently binds to a single complex non-canonical Retinoic Acid Response Element (RARE) in the LBR gene promoter (Schuler et al., 1994). Activation of LBR transcription appears to involve the interplay of numerous transcription factors (e.g., Sp1, AP-1, AP-2 and NF-kB).

A recent publication (Liu et al., 2023) studying mouse granulocytes, presented microscopic evidence that vimentin is responsible for nuclear “segmentation”. The authors claim that genetic deletion of vimentin eliminated the segmented nuclear shape, which then resembled granulocyte nuclei of human Pelger-Huet Anomaly. Our experiments with RA-induced HL-60/S4 granulocytes differ from their conclusions. As mentioned above (Figure 10), Log_2_FC of LBR is (1.56); but Log_2_FC of Vimentin is −4.88 and Plectin is −1.00. In our human myeloid HL-60/S4 cells, RA induced granulocytes exhibit nuclear lobules and ELCS, despite a considerable reduction of the VIM/PLEC cytoskeleton. We can only speculate that the species difference and/or different mouse granulocyte nuclear shape (i.e., donut or ring shape) might be the basis of their observations. We recommend future studies of a mouse LBR-mutated promyelocytic cell line “Ichthyosis” (*ic*), which undergoes RA-induced granulocyte nuclear differentiation from a nuclear chromatin sphere to a ring shape. Much has been described about these cells (Gaines et al., 2008; Shultz et al., 2003; Zwerger et al., 2008). It would be helpful if the transcriptomes for these cells before-and-after cell differentiation can become available.

## Supporting information

Table S1: Leading Edges of Cholesterol Biosynthesis Gene Sets.

## Acknowledgements

ALO and DEO express our gratitude to the MaineHealth Institute for Research for allowing us to join the research group of Dr. Igor Prudovsky. ALO and DEO thank Dr. Igor Prudovsky for opening his laboratory to us and for his exceptional generosity. Confocal microscopy studies were performed using a Leica SP-8 microscope at the Histopathology and Microscopy core (supported by NIH COBRE grant P30GM106391) of MaineHealth Institute for Research.

## Supplementary Materials

Table S1: Leading Edges of Cholesterol Biosynthesis Gene Sets.

